# Short-latency preference for faces in the primate superior colliculus

**DOI:** 10.1101/2023.09.06.556401

**Authors:** Gongchen Yu, Leor N. Katz, Christian Quaia, Adam Messinger, Richard J. Krauzlis

## Abstract

Face processing is fundamental to primates and has been extensively studied in higher-order visual cortex. Here we report that visual neurons in the midbrain superior colliculus (SC) display a preference for faces, that the preference emerges within 50ms of stimulus onset – well before “face patches” in visual cortex – and that this activity can distinguish faces from other visual objects with accuracies of ∼80%. This short-latency preference in SC depends on signals routed through early visual cortex, because inactivating the lateral geniculate nucleus, the key relay from retina to cortex, virtually eliminates visual responses in SC, including face-related activity. These results reveal an unexpected circuit in the primate visual system for rapidly detecting faces in the periphery, complementing the higher-order areas needed for recognizing individual faces.

**One-Sentence Summary:** An unexpected circuit through the primate midbrain reports the presence of a face in peripheral vision in 1/20^th^ of a second.

## Main Text

Faces are arguably the most important visual form stimulus for primates. Our ability to recognize and distinguish a remarkable number of faces and facial expressions, in a variety of contexts and from different viewpoints, involves the construction of invariant representations by complex visual circuits. This sophisticated processing of faces encompasses multiple regions of the cerebral cortex, most notably the “face patches” in the ventral visual pathway (*1-4*), as well as cortical regions in frontal cortex (*5, 6*) and subcortical regions such as the amygdala (*7-10*). Although recognizing the identity and expression of a face is a complex problem most commonly solved using foveal vision, detecting the presence of a face outside the fovea — regardless of its identity—is a simpler problem that could be accomplished by a basic circuit using conjunctions of visual features (*11, 12*). Such a rudimentary mechanism would be relatively coarse and less selective than those observed in the ventral visual pathway, but could have other advantages. It might be implemented at earlier stages of visual processing through early visual cortex, the visual thalamus, or directly from the retina (*13*), and therefore be substantially faster. If driven by coarser visual inputs at short latencies, it might be used to detect faces in the peripheral visual field (*14, 15*) that could then be summoned for fine-grained foveal analysis with a flick of the eyes.

Here we provide evidence that such a mechanism exists at the level of visual neurons in the primate superior colliculus (SC). We show that a preference for face stimuli at peripheral locations emerges rapidly in SC neurons, already within 50ms of stimulus onset, and provide causal evidence that this short-latency face preference depends on signals routed through early visual cortex.

### SC neurons exhibit a short-latency preference for faces

We recorded the activity of visually responsive neurons in the superficial and intermediate layers of SC in two macaques while presenting a variety of visual object stimuli at a peripheral (∼8 degrees eccentric) visual field location (Fig. 1A). The stimuli consisted of images from five object categories often used to assess category preferences in visual neurons: faces, bodies, hands, fruits/vegetables, and human-made objects (150 images overall, Fig. 1B).

**Fig. 1.**
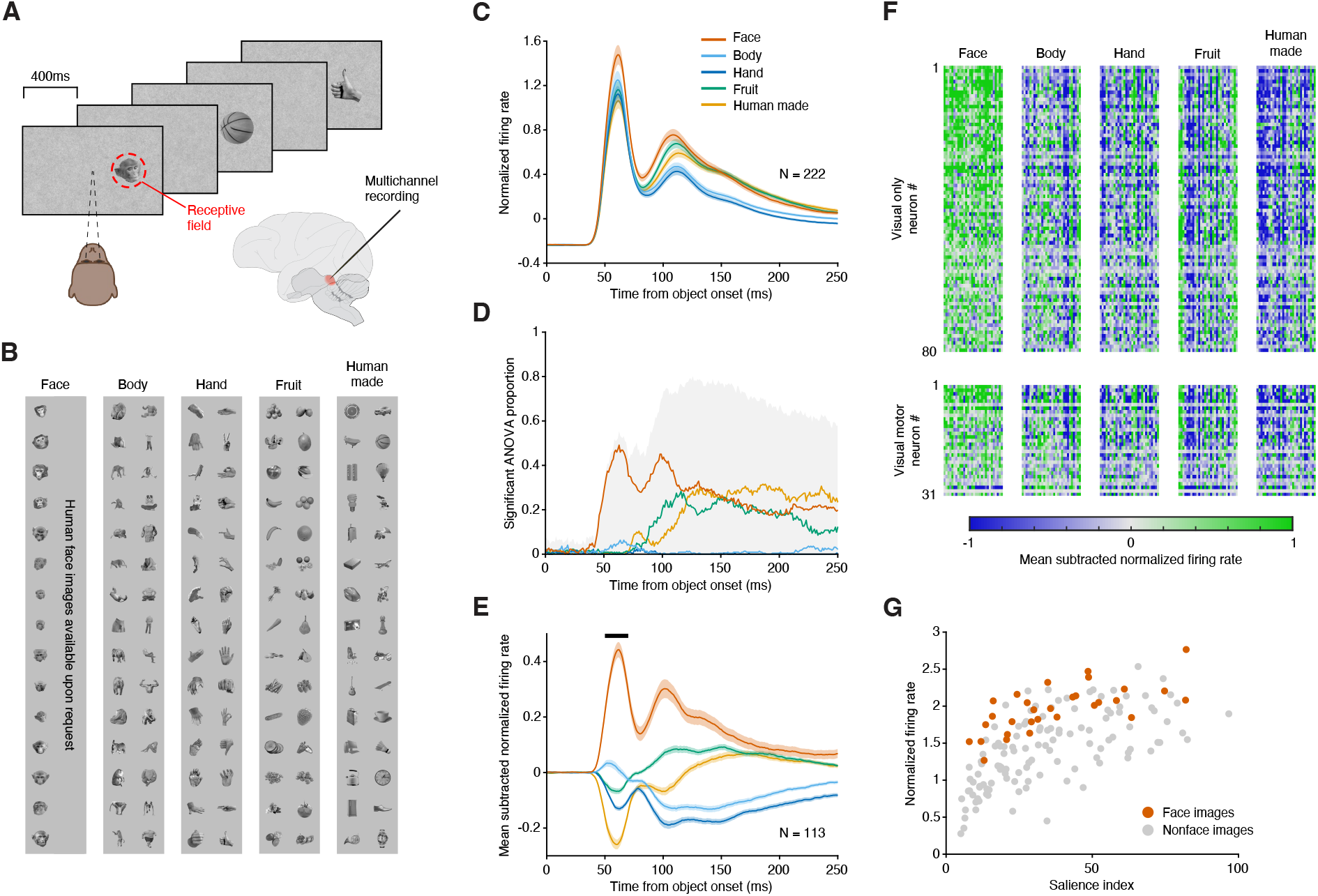
SC neurons exhibit a short-latency preference for faces. (A) Images were sequentially presented peripherally in the response field of the recorded SC neurons, while the monkey maintained central fixation. (B) The image-set consisted of 150 images, 30 each from five categories, matched for low-level features across categories. (C) Average of normalized firing rates over the population of 222 visually responsive SC neurons recorded in two monkeys, separately for each object category, aligned on image onset. (D) Proportion of SC neurons exhibiting object selectivity over time (ANOVA with category as group variable, p<0.05). Gray shaded area: overall proportion of object-selective neurons (preference for any object category). Individual curves: proportion of neurons displaying a preference for a specific object category. (E) Mean-subtracted normalized firing rates averaged over the SC neurons classified as object-selective in a 50-70ms time window (black bar). (F) Mean-subtracted average responses to each object image (columns, grouped by object category) for each neuron (rows), over the same time window. Upper boxes show data from ‘visual-only’ neurons, lower boxes show data from ‘visual-movement’ neurons. (G) Population average response to each image in the same time window as a function of visual salience. Red dots denote individual face images and gray dots denote individual non-face images. Error bars in panels C and E indicate the standard error of the mean (SEM).

To determine whether neurons in SC exhibit a true preference for a particular object category, it was crucial to account for variations in low-level visual features across stimuli. Otherwise, any observed category preferences might be trivially explained by a neuronal response preference for some low-level feature that was more prevalent in the images for one category, or even by one particularly salient image (*16, 17*). We therefore used a large number of exemplar images (30 per category) and matched the distributions of several low-level features (i.e., visual contrast, object size, and spatial frequency power in three distinct bands, see Methods, Fig. S1) across the five image categories. Using this carefully constructed image set we tested whether SC neurons exhibited any systematic preference for faces (or other object categories), independent of low-level visual factors.

Visually responsive neurons in SC responded most strongly to face stimuli compared to stimuli from all other categories. The population response of 222 SC neurons exhibited an early phasic response that started at ∼40ms and peaked approximately 60ms after image onset, followed by a second peak at 110ms (Fig. 1C). These temporal dynamics suggested two phases of object processing by SC neurons, which we verified by quantifying the prevalence of object preferences over time. In the early phasic response, and as early as 40ms after image onset, a large proportion of SC neurons exhibited a significant effect of object category (i.e., “object selectivity”, one-way ANOVA, p<0.05, gray shading in Fig. 1D). By the peak of this first phase (60ms after image onset), approximately half of the neurons exhibited object selectivity, with almost all of these neurons responding preferentially to faces (red curve in Fig. 1D). During the later phase of the visual response (>110ms after image onset), an even larger proportion of SC neurons exhibited object selectivity (close to 80%), but their preference was no longer overwhelmingly directed to faces, and instead was roughly equally split amongst faces, fruit and human-made objects. Thus, the initial visual response of SC neurons exhibited a strong preference for faces, and the latencies of this preference were extremely short, substantially shorter than those observed in object and face selective regions in temporal cortex where object selectivity typically emerges between 80 to 100ms following image onset (*1, 18*).

The short-latency preference for faces was not driven by particularly salient face images or by a small outlier group of face-preferring neurons. To document these points, we isolated the object-related visual modulation by taking the mean-subtracted response of our object-selective neurons to each of the 5 object categories; the population average showed a much stronger relative response to the face category (Fig. 1E). We next assessed the robustness of this effect across individual neurons and individual stimuli by examining the short-latency response (50-70ms window following image onset, black bar in Fig. 1E) of each neuron to all 150 exemplar images. The majority of object-selective SC neurons displayed stronger responses for most exemplars in the face category compared to exemplars in the other categories (Fig. 1F), and this held true for both classes of visually responsive neurons in our sample, functionally classified as “visual” and “visual-movement” (*19*).

The stronger response for face stimuli cannot be explained by differences in image saliency (*17*). First, by design, saliency did not differ between images of faces and images of non-face objects (Fig. 1G, KS test, p>0.05). Second, the preference for face stimuli was observed across a wide range of stimulus saliencies (ANCOVA, p<0.05). Finally, the early visual response increased with saliency (Fig. 1G), as expected (*17*), but the effects of salience and object category contributed independently to the amplitude of the response (ANCOVA, interaction, p=0.25).

We also verified that the preference for face stimuli was not due to other factors known to influence SC activity, including saccadic suppression or attention-related effects associated with microsaccades (*20, 21*) (Fig. S2) and visual adaptation (*22*) (Fig. S3). Lastly, while most of our neurons had peripheral receptive fields (∼8° eccentricity), in a separate set of experiments we studied SC neurons with foveal receptive fields and found the same results (Fig. S4). Thus, the early face preference we found in SC was driven by a variety of face stimuli regardless of their idiosyncratic low-level features, could not be trivially explained away due to saliency, microsaccades, or adaptation, and was present in both peripheral and foveal parts of the SC retinotopic map.

### SC neurons can be used to distinguish faces from non-faces with high accuracy

To assess the potential usefulness of this preference for face stimuli, we trained a series of binary classifiers based on the firing rates from our population of SC visually responsive neurons, and evaluated the cross-validated accuracy for each pair-wise classification of object categories (Fig. 2A, see Methods). Among all the binary classifiers, those that discriminated responses to faces from those to any other category were, already at the shortest latencies (within 50ms), the most accurate (Fig. 2A, green traces). In contrast, classifiers applied to pairs of non-face categories achieved lower accuracies, which improved somewhat over time (Fig. 2A, magenta and gray traces).

**Fig. 2.**
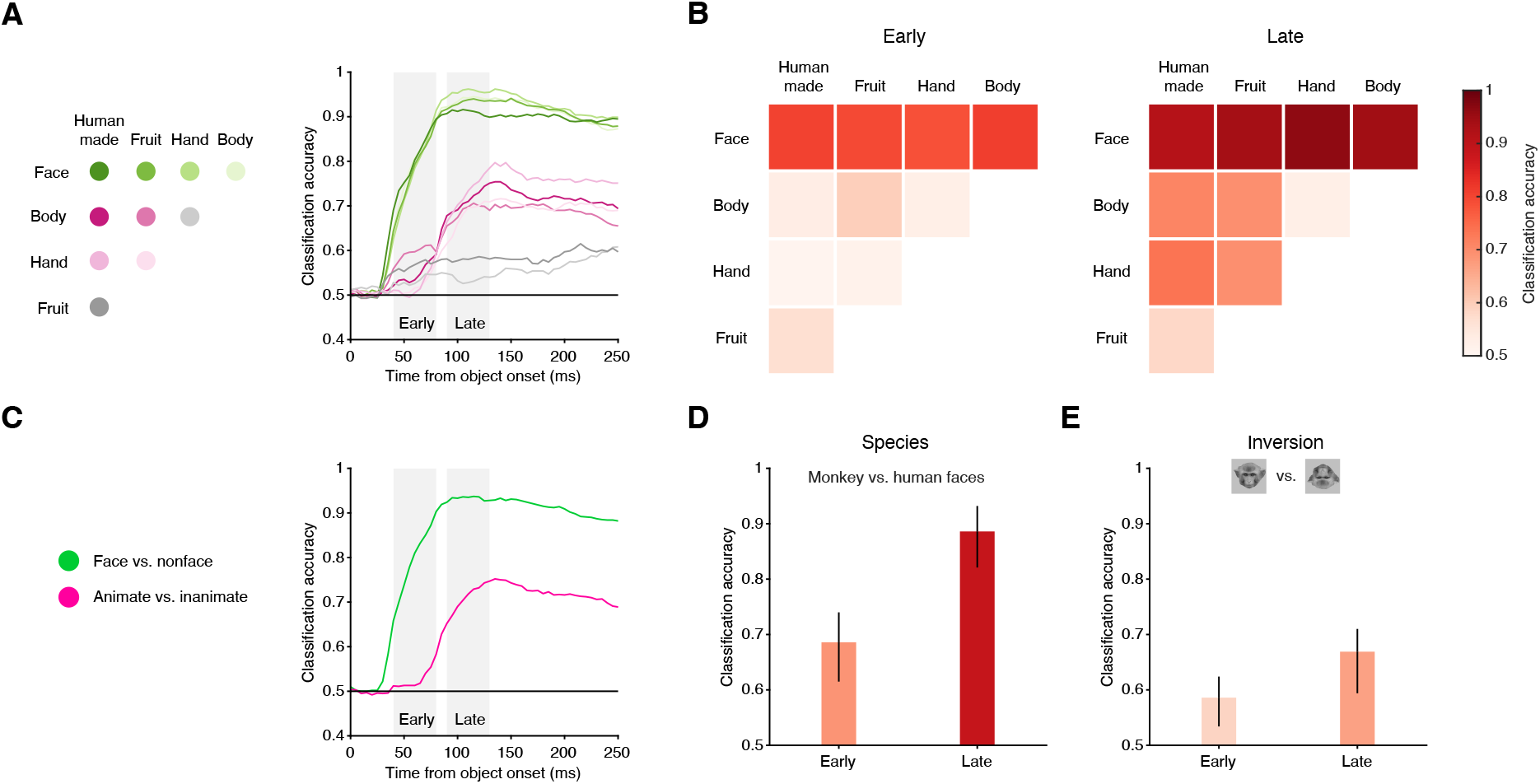
SC neurons can be used to distinguish faces from non-faces with high accuracy. (A) Left: legend denoting all possible pairwise relationships between individual object categories used for binary classification. Right: Cross-validated classification accuracy over time (aligned to image onset) for all binary classifiers trained on the binned (bin width 40ms) spike counts of SC visually responsive neurons to individual image presentations. Gray shaded rectangles indicate ‘early’ (40 to 80ms) and ‘late’ (90 to 130ms) windows used for summarizing classification accuracy in panels B and D. Note that classifiers were trained anew based on the spike counts in each bin. (B) Confusion matrices summarizing pairwise classifications of individual object categories in the ‘early’ (left panel) and ‘late’ (right panel) windows. (C) Left: colors denote grouping of object categories into pairwise classifiers for images that contain faces versus non-faces (i.e., faces vs. human made, fruit, hand, and body, in green) and images that contain animate objects versus inanimate (hand and body vs. human made and fruit, in purple). Right: Cross-validated classification accuracy over time for binary classifiers on these groupings. (D) Accuracy for classifying species (i.e., images that contain monkey faces vs. human faces) in the ‘early’ and ‘late’ windows. Image icons are presented as an example. Error bars denote 95% confidence interval, bootstrapped. (E) Accuracy for classifying image inversion (i.e., upright vs. inverted faces) in the ‘early’ and ‘late’ windows. Same format as (D).

To summarize these results, we focused on two temporal windows, one early (40 to 80ms after image onset) and one late (90 to 130ms), that encompassed the two peaks in the population response (Fig. 1C). For each window, we constructed a confusion matrix to indicate the accuracy for each pairwise object-category classification (Fig. 2B). In the early window, faces were the only category that could be accurately distinguished from other objects. During the late window, classification of faces further improved. Furthermore, animate objects (bodies and hands) could now be distinguished from inanimate objects (fruits/vegetables and human-made objects) at reasonable accuracies. Note that our ‘animate’ stimuli did not include any images of faces.

Pooling the results into ‘face vs. non-face” and “animate vs. inanimate (non-face)” further highlights the differences in the timing of object category preferences in SC (Fig. 2C). Faces could be discriminated from non-faces with accuracies that were significantly above chance at 40ms after image onset (95% confidence interval above chance level of 0.5), reached 80% accuracy by 60ms, and peaked at an accuracy of 92% at 100ms (Fig. 2C, green). In contrast, the ability to accurately classify animate from inanimate images (Fig. 2C, magenta) emerged later (at ∼75ms) and was weaker (peaked at 75%). Thus, the preference of SC visual neurons for object categories was almost exclusively related to faces at short latencies, and included other object categories only later, possibly due to the later arrival of object-related signals from higher-order cortical areas.

In the cortical face-patch system, neurons respond more strongly to faces of conspecifics compared to faces of a different species and prefer upright over inverted faces (*1, 23, 24*). Such neuronal preferences may underlie the psychophysical findings that primates are better at detecting faces of conspecifics (compared to faces of other species), as well as at detecting upright faces (compared to inverted faces) (*25, 26*). We found a similar effect in SC. Based on our SC visual responses, we found that images of monkey faces could be accurately distinguished from images of human faces (Fig. 2D) already in the early time window (69%), and improved in the later window (89%). Additionally, upright faces could be accurately distinguished from inverted faces (Fig. 2E) in the early window (59%) as well as in the later window (67%). Thus, the face preference of visual neurons in primate SC also has some of the selective properties observed in classic face-selective cortical areas, although more so at longer latencies.

### Short-latency face preference in SC depends on visual cortex

We next investigated the circuit underlying this short-latency face preference. The two primary sources of visual inputs to the SC are the retina (*27*) and, by way of the lateral geniculate nucleus (LGN), the primary visual cortex (V1) (*28*). Direct retinal input to SC have been put forward as a fast subcortical route for processing emotionally relevant events (*29*) and proposed to play a key role in fast face detection (*30*). We reasoned that if the face preference in SC is driven by retinotectal (as opposed to corticotectal) projections, then disrupting the corticotectal pathway would leave intact the short-latency face preferences in SC. To test this idea, we again recorded in SC while presenting face and non-face stimuli (hands and human-made objects only in these experiments) at a peripheral visual field location and compared the degree of face preference in SC before versus during temporary inactivation of LGN via muscimol (Fig. 3A).

**Fig. 3.**
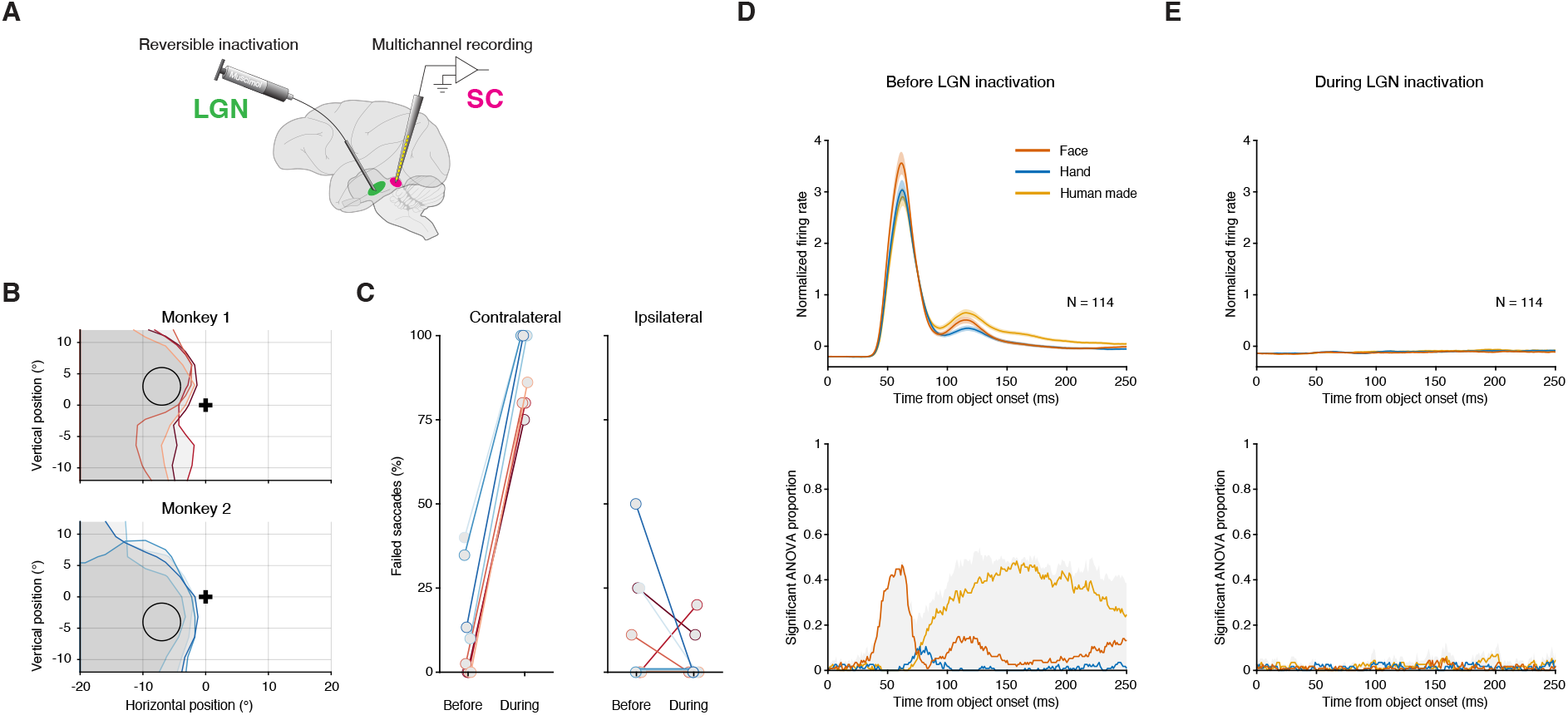
Short-latency face preference in SC depends on visual cortex. (A) SC neurons were recorded before and during reversible inactivation of LGN by local injection of muscimol. (B) Maps of the visual field region to which monkeys failed to complete a visually guided saccade (scotoma). Scotomas for each session are plotted, separately for each monkey. Black cross: fixation spot; circle: mean RF location of SC neurons across sessions, where the object stimulus was placed; gray shaded areas: scotoma region on each experimental session (different contour colors). (C) Percentage of saccade failures into the RF of SC neurons before and during LGN inactivation (contralateral to the manipulated LGN, left panel) and to a diametrically opposite control location (ipsilateral, right panel). Each connected pair of circles reflects data from one experimental session. (D) Average normalized firing rates over the population of visually responsive SC neurons recorded before LGN inactivation (top panel) and proportion of such neurons exhibiting object selectivity (bottom), aligned to image onset. Similar format to figure 1 C and D. (E) Same as D, over the same set of neurons, but during LGN inactivation. Error bars denote SEM.

LGN inactivation induced a large, but retinotopically confined, “scotoma” – i.e., an area in the visual field where monkeys could no longer report the occurrence of visual events, consistent with previous results (*31*). We measured the extent of the scotoma by tabulating the range of locations in the visual field associated with failures on a simple visually guided saccade task (Fig. S5) and verified that the affected region encompassed the SC neurons’ receptive fields, which is where the task stimulus was positioned (Fig. 3B). The percent of failed saccades into the stimulus area in the hemifield contralateral to the manipulated LGN increased from 13% before LGN inactivation to 90% during (average over sessions, p < 0.01, Wilcoxon Sign test, Fig 3C). Saccades to the ipsilateral hemifield were unaffected by the manipulation (p = 0.31).

Before LGN inactivation, we replicated our findings from Figure 1 in a population of 114 SC neurons including category selectivity that was almost exclusively driven by a preference for faces at short latency, and a mix of category preferences at longer latencies (Fig. 3D).

During LGN inactivation, visual responses in these same SC neurons were largely abolished, including the earliest phasic response (Fig. 3E). This loss of responsiveness was specific to visual stimuli and not a general reduction in neuronal excitability, because SC neurons that were recorded continuously throughout the experiment continued to exhibit normal levels of activity during LGN inactivation in a control period that did not include visual stimuli (Fig. S6).

This elimination of visual responsiveness was observed for all object stimuli regardless of content and category. The proportion of neurons exhibiting object selectivity did not exceed chance levels throughout the presentation of the visual stimulus, and a preference for faces was never observed (Fig. 3E). Thus, contrary to expectations of a fast subcortical route directly from the retina to the SC, the face preference, and in fact, all the visual responses we measured, depended crucially on signals mediated through the LGN. There are no known projections from LGN to the SC in the primate, but short-latency visual signals are prominently relayed from LGN to V1 and then to the SC. Our LGN inactivation findings, therefore, point to a visual circuit involving V1 to the SC as the most likely source of the short-latency face preference we observed.

Several additional findings support our interpretation that our causal manipulation was specific to LGN-mediated inputs to visual cortex as opposed to other inputs. First, we confirmed the placement of our muscimol injection site to the peripheral visual field representation of LGN by tracer injection and histology (Fig. S7). Second, muscimol does not affect fibers of passage (*32*) so the effects could not be attributed to silencing either the optic tract or retinotectal projections that pass nearby. Third, in a set of additional control experiments we confirmed that retinotopic overlap between the visual scotoma and receptive fields in the SC was necessary to observe the effects (Fig S8), so structures adjacent to the LGN (that lack retinotopic structure) could not be the cause of the effects. Thus, the loss of visual responses in SC neurons during LGN inactivation was due to a specific disruption of the pathway leading from geniculate to the SC, through visual cortex.

### Plausibility of a corticotectal circuit for early face processing

We next determined whether it was plausible that the outputs of V1 could provide SC with the necessary signals to construct a preference for faces at short latencies (Fig. 4). We constructed a multi-scale computational model based on V1 simple cells comprised of Gabor filters that varied in RF location, orientation, spatial frequency, and phase (see Methods). The filters and their readout weights were essentially a two-layer network that emulated a very simple version of visual processing in V1 that might provide inputs to visual neurons in the SC. We then applied the same cross-validated binary classifier analysis we used on our SC data to the outputs of this V1 model to our stimuli, and evaluated whether face images could be discriminated from non-face stimuli based on these V1 model responses.

**Fig. 4.**
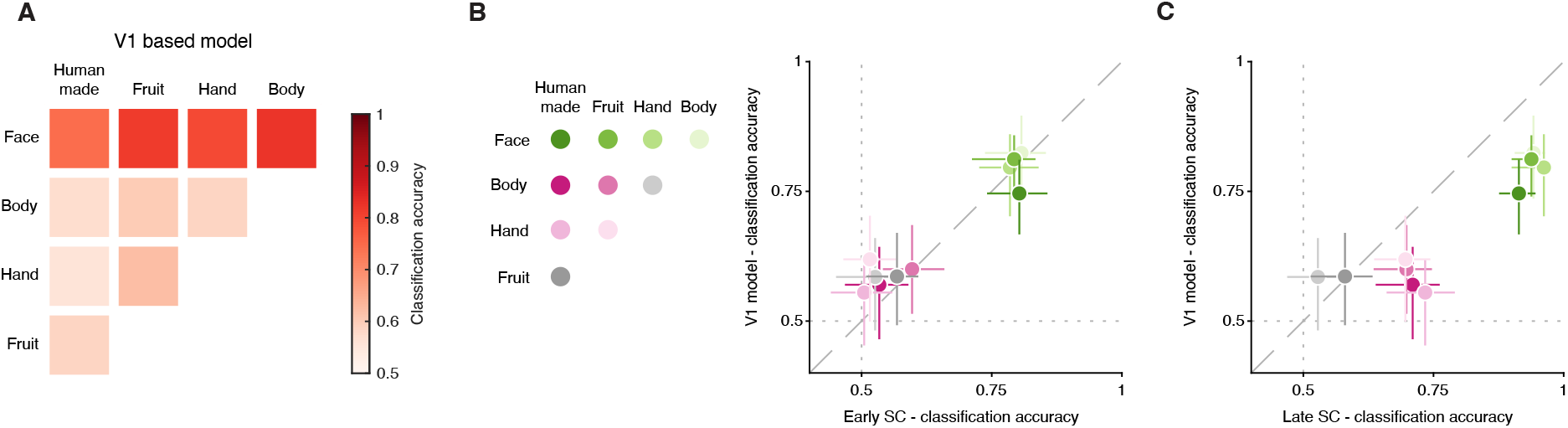
Plausibility of a corticotectal circuit for early face processing. (A) Confusion matrix summarizing the cross-validated performance of binary classifiers trained on the outputs of V1-like model cells to the images in our dataset. (B) Comparison between the classification accuracy of binary classifiers trained on V1-like model cells outputs (ordinate) or on spike counts emitted by our 222 SC neurons in the early (40 to 80ms) time window. One data point for each binary classifier (color-coded, see legend). (C) Same as B, but for SC spike counts in the late (90 to 130ms) time window. Error bars denote 95% confidence interval, bootstrapped.

The results were remarkably similar to those obtained with our SC neuronal data. Face images could be classified from non-face images at high cross-validated accuracies of ∼80%, regardless of which other non-face category was tested (Fig. 4A, top row), whereas other pairwise classifications produced poor performance (Fig. 4A, bottom 3 rows). These model-based results are on par with the classification performance we observed with our SC neuronal data, but only when compared to the classifier results in the early window (Fig. 2B), consistent with the latency of presumed V1 inputs to the SC. Direct comparison of the results based on the V1 model and our SC data confirms that classifier performance in the early visual response (40–80ms after image onset) is consistent with input signals arriving from V1 (Fig. 4B) whereas performance in the late epoch (90–130ms) is not (Fig. 4C). Applying the same classifier analysis to the output of an LGN-based model (see Methods) produced results that matched neither the early nor the late SC data, yielding much poorer performance (Fig. S9). These results show that a visual circuit from V1 to the SC could plausibly generate the short-latency face preference we observed. In contrast, the object preferences observed later in the SC (after 100ms) could not be recapitulated by our V1 model and presumably depend on inputs from other, higher-order visual areas.

## Discussion

Our results provide new insights into how primates process faces. We have identified a circuit operating at an early stage of the visual system—the midbrain superior colliculus—for detecting faces in under 50ms. Causal manipulations revealed that the visual responses to face stimuli and in fact, to all object stimuli, depend on signals routed through early visual cortex. The function of this midbrain-centered mechanism seems complementary to the more advanced processing centered in temporal cortex, that supports the recognition of individual faces and facial expressions across a range of viewpoints and contexts (*33*). By detecting faces rapidly, and operating at peripheral as well as foveal parts of the visual field, this midbrain circuit may play a crucial role in detecting faces and then promptly triggering the targeting eye movements (*34, 35*) that bring the face to the fovea for finer-grained analysis by higher-order cortex. The short-latency face preference reported here might therefore explain the propensity of primates to orient towards faces at extremely short latencies (*36*). Our study was done in adult macaques, but if similar properties were present at birth, this could explain the tendency of newborns to preferentially fixate on faces, despite having yet to develop cortical face patches (*4, 37, 38*). Since the maturation of visual cortex depends on visual experience (*39, 40*), disruptions in this midbrain circuit during development might interfere with the normal development of face patches and other higher-order functions important for social interactions (*41-43*).

## Acknowledgments

We thank Nick Nichols, Daniel Yochelson, Denise Parker and Hayden Warnock for technical support. We thank Carlos Mejias-Aponte and Martin Bohlen for assistance with the tracer injection and image processing. We are grateful to David Leopold and Arash Afraz for providing feedback on an earlier version of the manuscript, and to Kara Cover, Divya Subramanian, Xuefei Yu, Marianne Duyck, Stuart Duffield, Kerry McAlonan and James Cavanaugh for helpful discussions. This work was supported by the National Eye Institute Intramural Research Program at the National Institutes of Health (ZIA EY000511).

## Author contributions

Conceptualization: GY, LNK, RJK

Methodology: GY, LNK, CQ, AM

Investigation: GY, LNK, CQ

Visualization: GY, LNK, CQ

Funding acquisition: RJK

Project administration: RJK

Supervision: RJK

Writing – original draft: GY, LNK, RJK

Writing – review & editing: GY, LNK, CQ, AM, RJK

## Supplementary Materials

Materials and Methods

Figs. S1 to S10

