## Supplementary materials for "Short-latency preference for faces in the primate superior colliculus"

**The PDF file includes:**

Materials and Methods
Figs. S1 to S10

### Materials and Methods

General

All experimental protocols (protocol number #NEI-649) were approved by the National Eye Institute Animal Care and Use Committee, and all procedures were performed in accordance with the United States Public Health Service policy on the humane care and use of laboratory animals. Data collection and analysis were not performed blind to the conditions of the experiments.

Subjects

We collected and analyzed data from two adult male rhesus monkeys (Macaca Mulatta, aged 13 to 16 years, weight 9 to 12 kg). A plastic headpost and recording chamber had been previously implanted granting electrophysiological access to both the superior colliculus (SC) and lateral geniculate nucleus (LGN).

Behavioral apparatus

Monkey subjects were seated in a customized primate chair (Crist Instrument Co., Hagerstown, MD, United States) and head-fixed inside a darkened booth, in front of a VIEWPixx display (VPixx Technologies, refresh-rate 100Hz). The progression of the experiments was orchestrated using a modified version of PLDAPS (*1*). Subjects' real-time horizontal and vertical eye position data were recorded at 1000Hz using an EyeLink 1000 infrared eye-tracking system (SR Research Ltd.). Trials were initiated by a manual joystick press, collected with a single-axis joystick (CH Products, model HFX-10) mounted to the primate chair and oriented to allow vertical movement.

Saccade task

Prior to running the object viewing task detailed below, monkeys performed a standard visually guided saccade task to map the SC neurons' receptive field (RF) location and extent. A memory guided version of the task was used to determine the functional class of each neuron. Additional details have been published elsewhere (*2*).

Object viewing task

Following RF mapping using the saccade task, subjects performed an object viewing task (Fig 1A). Trials were initiated when the monkey pressed down a joystick to trigger the appearance of a 0.25° wide white fixation square (48 cd/m$^2$) on a gray background (32.6 cd/m$^2$). Fixation had to be maintained within a 1.5° wide square window (invisible to the monkey). After acquiring fixation, the gray background was replaced with a pink noise texture with the same average luminance (RMS contrast of 4.38%) and following a 500ms delay, 3 to 5 visual objects were sequentially presented at a prescribed location (3 - 9° eccentricity depending on the receptive field locations of the recorded neurons). Each object was presented for 400ms and was separated from the presentation of the next object by a 400ms interval. On every object onset and offset, the pink noise background was refreshed to a new texture at random. Maintaining fixation throughout the entire course of a trial netted the monkeys a liquid reward.

Image set

We used a total of 150 grayscale images of objects. Object images were selected from five object categories: faces (30 images), bodies (30 images), hands (30 images), fruits and vegetables (30 images), and human-made objects (30 images) (Fig. 1A). In addition to the 150 images from the five categories, we included three vertical Gabor patches with a carrier spatial frequency of 0.33, 1.66 and 5 cycles/degree, and one image of a solid gray patch. Within the faces, bodies, and hands category, 15 of the 30 images were from humans and 15 were from monkeys. To determine whether SC neurons exhibit a preference for one or more categories over others it was important to account for variations in image low-level features across categories. Otherwise, a category preference could trivially emerge (e.g., if images in a category had higher average contrast). We thus manipulated our original images (using the SHINE toolbox (*3*) and custom code) to match the luminance, RMS contrast, image size, and power in low, medium and high spatial frequency bands across categories (for precise values ranges see Fig. S1).

Matching the low-level image features already allowed us to interpret any differential response in SC neurons as driven by the image category, and not by image features. However, some variation in low-level features still exists across images. To account for their potential effect on SC neurons' responses we constructed a multilinear regression model (Fig. S10). We regressed out the effect of low-level features on SC responses and evaluated the effect of object category on the residuals. Results were consistent with those present in the main text, albeit larger in magnitude. For the analysis presented in Figure 1G, salience was computed using the GBVS algorithm (4). Because this model only computes a salience map of a visual scene, we randomly arranged all our images on a grid, and then measured, for each of our stimuli, the average salience within the region of the image in which it was placed. Because relative location of the images can affect the saliency map, we repeated this process 100 times and averaged the results.

SC recordings

Electrophysiological data were acquired by an Omniplex-D system (Plexon Inc.). To record neuronal activity in SC, 32-channel Plexon v-probes with 50 μm inter-channel spacing (Plexon Inc.) were used. A motorized microdrive (NAN Instruments) was used to carefully advance the probes to their target depth, aiming for the superficial and intermediate layers of SC. Target depth was based on functional properties of the recorded neurons (i.e., functional class): we aimed to record visual neurons (consistent with the superficial layer of SC) as well as visual-movement neurons (consistent with the intermediate layers). We then adjusted probe position to maximize neuronal yield and left it to settle for ~1 h before starting the experimental session, to stabilize the tissue and improve recording quality. Given the angle of approach of the recording probe, most neuronal RFs overlapped. We placed task stimuli in a location that maximized overlap across RFs.

LGN inactivation

LGN was approached through the same chamber used to access the SC. To access both structures at the same time we used a hybrid grid (the "hygrid", made in house) with two sections: one consisting of straight grid holes leading directly to the SC and a second with angled grid holes leading to the LGN. The LGN was approached based on MRI scans and identified by observing strong visually evoked activity consistent with LGN laminae and retinotopy (*5*). Reversible inactivation of the LGN was performed by injecting muscimol (0.8-1.2 µl; 5 mg/ml) into a custom made injectrode using a syringe pump (Legato, KD Scientific) at a constant rate of 0.1 µl/min. Thirty minutes after the end of muscimol injection, we used a visually guided saccade task to confirm that inactivation caused a scotoma, i.e. an area in the visual field over which visual events were no longer registered, and the monkey failed to saccade to (*6*). On each of the 8 sessions in which SC was recorded from both before and during LGN inactivation, we verified that the extent of the scotoma overlapped with the stimulus location, which matched the location of SC RFs. If an overlap was not achieved, we either injected more muscimol and/or waited an additional 20 minutes.

The monkeys' inability to register visual events in the scotoma is presented both qualitatively (Fig. 3B) and quantitatively (Fig. 3C). For the data in Fig. 3B, visual space was binned into 3.2° square bins and the mean failure rate to saccade was computed. These values were smoothed and the $40^{th}$ percentile contour plotted. For the data in Fig. 3C, failure rate was computed for saccades into the circular aperture in which the stimulus was presented (3° radius, indicated in Fig. 3B), contralateral to the manipulated LGN, before versus during LGN inactivation. For the ipsilateral hemifield a larger area was used (6° radius, mirror-opposite coordinates) due to lower spatial sampling in that hemifield.

To ensure proper fixation, we aimed to keep the scotoma from involving the foveal region. We therefore aimed to inject at the rostral pole of the LGN, an area of the nucleus that corresponds to the peripheral visual field (*5*).

On monkey 1's final experimental session, a bidirectional tracer 5% fluor-emerald (D3306, Invitrogen) was mixed with muscimol and injected into the same LGN location as in our previous inactivation sessions. The muscimol's effect on saccadic behavior was used to confirm successful injection; the tracer was used to confirm our anatomical location within the LGN. Four days after the injection, the brain was obtained. Fluorescent digital photomicrographs of histological sections were acquired using a Zeiss axio scan (Zeiss) and confirmed that our injection site hit the rostral pole of the LGN (Fig. S7), consistent with our expectation given the peripheral location of the induced scotoma.

Electrophysiological analysis

Only data recorded during successfully completed trials were used for analysis. Continuous spike-channel data collected during the experimental session were sorted offline with Kilosort2 (*7*) using in house tools (https://github.com/ElKatz/kilo2Tools), and manually curated by a human expert using Phy2 to ensure that all sorted units have plausible inter-spike interval distributions and waveform shapes consistent with action potentials. Neurons were excluded from analysis if they had low (<1.8) signal-to-noise ratio (*8*), low average trial firing rate (<1 spikes/s), or no clear visual RF. Overall, 401 neurons were recorded (221 from Monkey #1, 180 from Monkey #2) and 179 of these did not meet our inclusion criteria, netting 222 neurons for the analysis reported here (117 from Monkey #1, 105 from Monkey #2). Results did not differ across monkeys and were therefore combined to increase statistical power.

For visualization of mean firing rates over time relative to key events in the task (Fig. 1C), spike counts were binned into overlapping 20ms bins sliding in 1ms step. Each neuron's data was z-score normalized by subtracting the mean and dividing by the standard deviation across binned spike counts across trials and conditions.

Object selectivity was computed for each neuron over time (Fig. 1D) by performing a one-way ANOVA on spike counts for all 150 object images (total d.o.f. = 149, error d.o.f. = 145) with object category as the main factor (d.o.f. = 4), in overlapping 20ms bins sliding in 1ms step, aligned to stimulus onset. A neuron was considered object selective for $p < 0.05$. The object category which induced the largest neuronal responses was defined as the preferred category of the neurons.

Binary decoders for image category classification

We used binary linear classifiers to determine whether the responses of SC neurons could be used to correctly classify the image being presented to the animal. We considered comparisons between all pairs of the five classes presented to the animal, each having 30 members; we also tested the ability to discriminate faces (30 images) from all other stimuli (non-face, 120 images). In all cases we randomly split the images in each class in two equal-sized groups and used one group for training the classifier and the other for testing it. All performance measures reported refer to the performance on testing images. The response of each cell to each presentation of an image was computed by counting the spikes emitted by the cell in a 40ms window. For each cell we then sampled, with replacement, 1000 trials from those in which images from each of the two classes under consideration were presented, separately for training and testing image sets (4000 trials overall). Classification was then performed using all the neurons recorded, across all sessions and in both monkeys. Since the neurons were not all recorded simultaneously, the analysis is thus based on pseudotrials (and thus ignoring information potentially carried by noise correlations among neurons). A support vector machine with a linear kernel was then trained using the 2000 training pseudotrials, and its performance was evaluated on the 2000 testing pseudotrials. Accuracy was computed as the fraction of correctly classified testing images. This process was repeated 100 times, and median and 95% confidence intervals of the performance were reported. To evaluate classification performance over time, spike counts were counted in overlapping 40ms windows, sliding in 5ms steps. Independent classifiers were then trained and tested for each time bin.

V1-based model

To ascertain which signals related to our image set might be made available by V1 to the SC, we implemented a multi-scale model of V1 simple cells and applied linear classification analysis (using the same procedure as for our SC recordings) to the outputs of the V1 model. Our model used Gabor filters, followed by a half-rectifying nonlinearity (negative values are set to zero, positive values are kept unchanged), the standard for models of V1. We considered four scales, with wavelengths of 6 deg (the size of our images), 3 deg, 1.5 deg, and 0.75 deg. The standard deviation of the Gaussian envelope of the Gabor filter was set at 40% of the wavelength, as observed on average in macaque V1 (*9*). We did not consider finer scales for two reasons: First, our images were typically presented at ~8 deg of eccentricity, where higher spatial frequencies are not resolved (*10*); Second, to account for imperfect fixation, when we simulated the responses of these filters to our images, we jittered the images by up to 1 deg in each direction (with vertical and horizontal displacements independently drawn from random uniform distributions). The output of smaller scales would be completely confounded by such jittering. At each scale we then considered Gabor filters of 4 different orientations (spaced 45 deg apart) and 4 different spatial phases (spaced 90 deg apart). The filters tiled the image space, with the largest scale extending outside it. Overall, our model was composed of 7712 simulated V1 simple cells, whose outputs were then fed to the linear classifier.

<u>LGN-based model</u>

While LGN is not known to project to the SC in primates, we also implemented a simple model of LGN's on- and off-center circular symmetric cells. For this model we also considered four scales, which were selected to cover the same range of preferred spatial frequencies as the V1 model. At each scale the LGN filter was a difference of circular Gaussian functions, followed by a half-rectifying nonlinearity, the standard for models of LGN. The standard deviation of the center Gaussian was set to one eight of the wavelengths used in the V1 model, and that of the surround Gaussian was 5 times as large. We considered two phases for the cells, 0 (positive center, negative surround) and 180 deg (negative center, positive surround). Overall, our model was composed of 964 cells, whose outputs were then fed to the linear classifier.

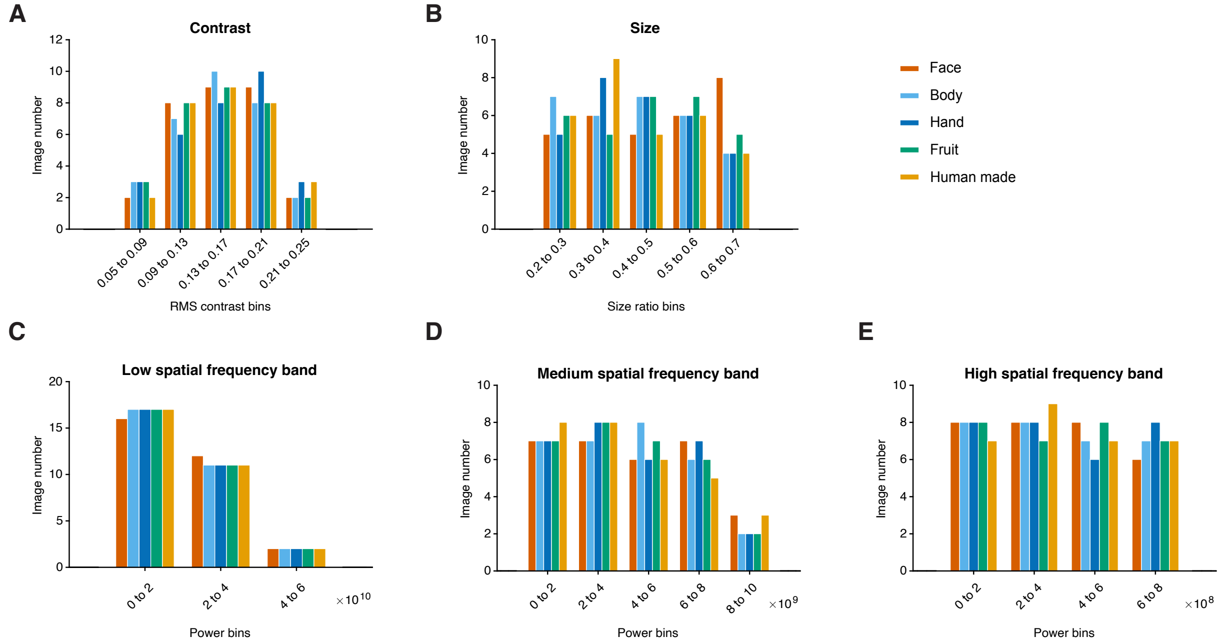


**Fig. S1. Distributions of low-level visual features were matched across object categories.** Five low-level features were selected, and the distribution of each was matched across the five object categories. The features are: (A) RMS contrast; (B) Size ratio (number of object pixels divided by number of pixels in the 6 deg aperture); (C) Spectral power in a low spatial frequency (SF) band (0.1667 to 0.6525 cycles/degree); (D) Spectral power in an intermediate SF band (0.6525 to 2.5544 cycles/degree); (E) Spectral power in a high SF band (2.5544 to 10 cycles/degree). Distribution matching was confirmed with the Kolmogorov–Smirnov test ($p > 0.05$, for each of the possible pairwise comparisons of object categories for each low-level feature). Additionally, mean luminance of all object images was matched to the mean luminance of the background (32.6cd/$m^2$).

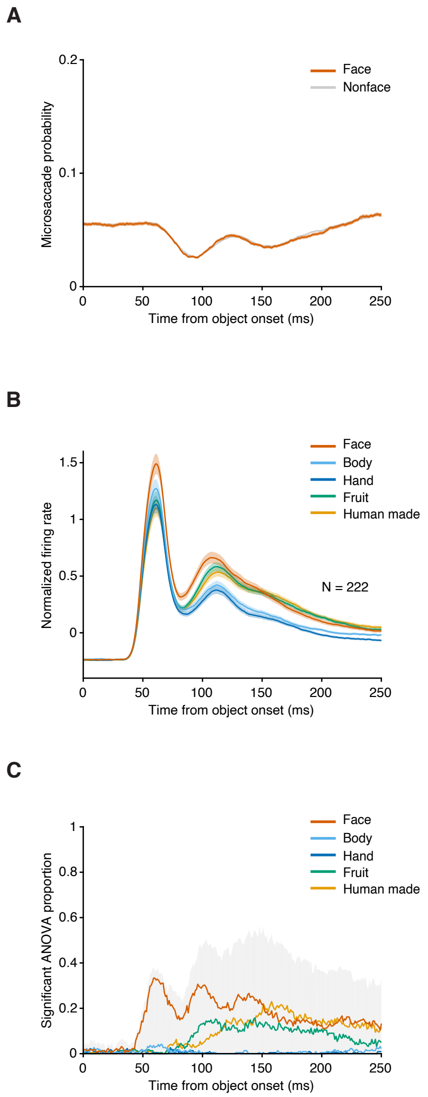


**Fig. S2. Short-latency face preference in SC cannot be explained by the presence of microsaccades.**

(A) Probability of microsaccades over time in 20ms non-overlapping bins aligned to image onset, for images from the face category (red) versus all others (gray). There is no significant difference in microsaccade probability between the two conditions, in any of the bins (Chi-square $p > 0.05$). (B) Average normalized firing rates over our population of visually responsive neurons when using only epochs during which no microsaccades occurred from 400 ms before image onset to 100ms after. Averages computed separately for each of the five object categories, aligned to image onset. (C) Proportion of SC neurons exhibiting object selectivity using the same epochs (ANOVA with object category as group variable, $p<0.05$). Gray shaded area: overall proportion of object-selective neurons (preference for any category). Individual curves: proportion of neurons displaying preference for a specific object category.

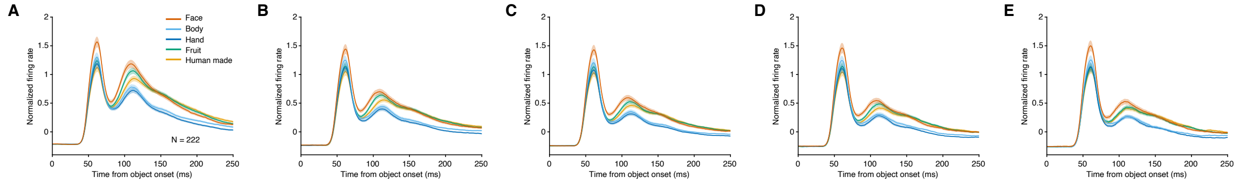


**Fig. S3. Short-latency face preference in SC cannot be explained by visual adaptation.** Average normalized firing rates over our population of SC visually responsive neurons across the five object categories, split by stimulus sequence order. Panel A shows data for images that appeared $1^{st}$ within a trial's sequence of images, panel B for images that appeared $2^{nd}$, and so on. Face preference at short latencies was observed in all cases.

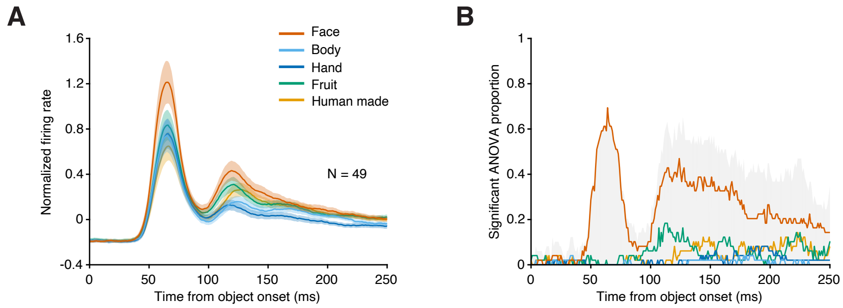


**Fig. S4. Short-latency face preference is also observed in SC neurons with foveal receptive fields (rostral SC).**

(A) Average normalized firing rates over a population of 49 visually responsive neurons recorded in rostral SC with foveal receptive field (eccentricities up to 2.7°), separately for each of the five object categories. (B) Proportion of SC neurons exhibiting object selectivity (ANOVA with object category as group variable, $p<0.05$). Gray shaded area: overall proportion of object-selective neurons (preference for any category). Individual curves: proportion of neurons displaying preference for a specific object category. Error bars denote SEM.

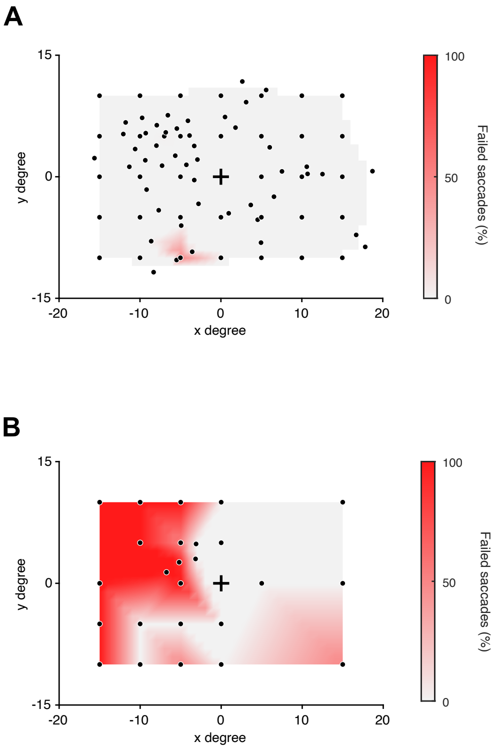


**Fig. S5. LGN inactivation induced a visual scotoma in a visually guided saccade task.**
Failure rate of saccades towards visual targets before (A) and during (B) LGN inactivation in one example session. Black cross: fixation location: Individual black dots: saccade target locations. (A) Before LGN inactivation, the failure rate is 0% for most targets in the visual field. (B) During LGN inactivation, there is a high failure rate for targets appearing in the top left corner of the visual field, indicating the scotoma location. Heatmaps are interpolated between target locations.

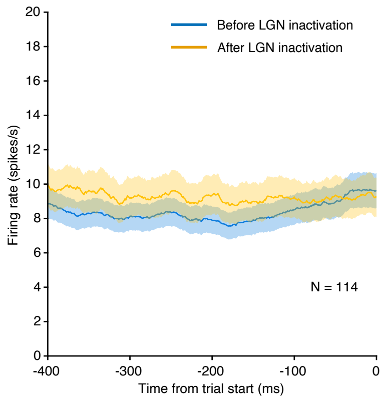


**Fig. S6. LGN inactivation did not reduce SC response during epochs with no visual stimulation.**

Average firing rates over our population of SC visually responsive neurons during an epoch that did not include visual stimulation: -400 to 0ms relative to the trial start, before (blue) and during (orange) LGN inactivation. Error bars denote SEM.

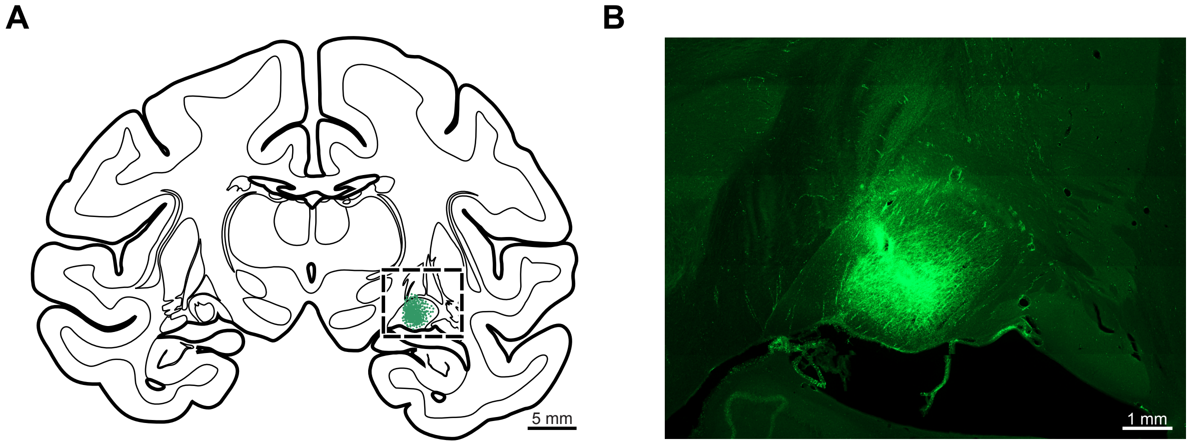


**Fig. S7. Histological confirmation of LGN injection site.**

(A) Low magnification camera lucida reconstruction of a 75 µm thick coronal section containing rostral LGN. Green stippling: approximate location and spread of the tracer infused into the right LGN. Box: approximate location of the photomicrograph shown in B. (B) Fluorescent digital photomicrograph illustrating a higher magnification view of the injection location and spread within the rostral LGN.

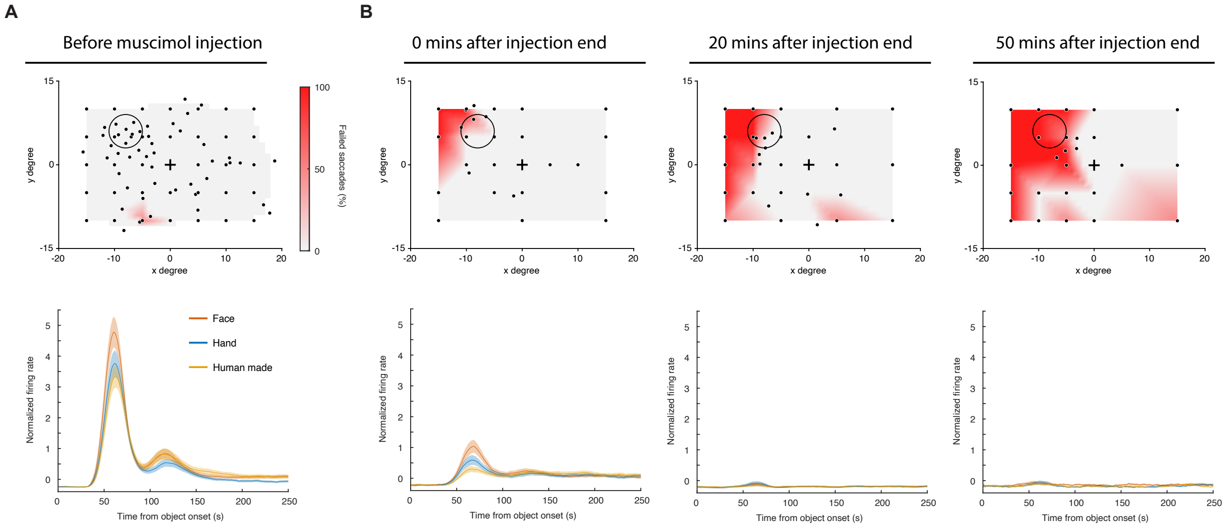


**Fig. S8. Retinotopic overlap between visual scotoma and SC receptive fields was necessary to observe the reduction in SC visual responses.**

Evolution over time of visual scotoma and SC responses to object images during an example LGN inactivation session. (A) Top: Failure rate for saccades towards visual targets prior to muscimol injection in LGN (similar format to Supplementary figure 5, but with the addition of a 6° diameter circle denoting the stimulus location used in the object viewing task, positioned to maximally overlap with recorded neurons' receptive fields). Saccade failure rate was 0% for almost all targets. Bottom: Average normalized firing rates over our population of SC visually responsive neurons prior to muscimol injection in LGN (similar to Figure 3D). (B) Same as (A), following injection of muscimol in LGN.

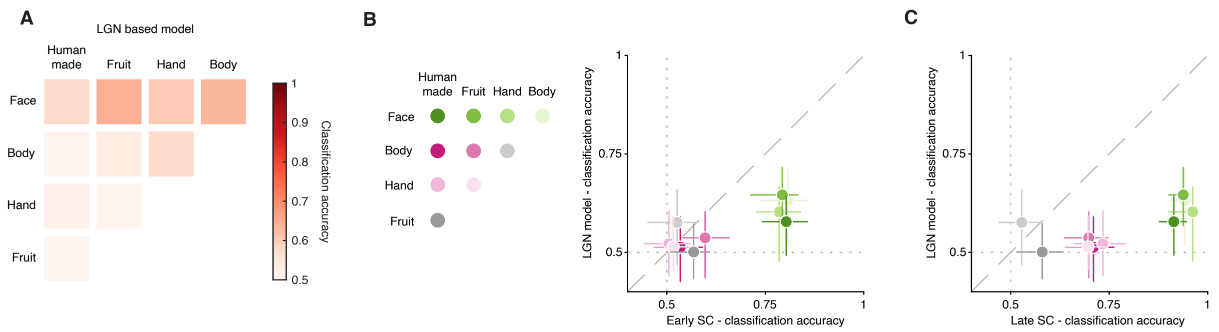


**Fig. S9. A LGN-based model matched neither the early nor the late responses of SC neurons.**
(A) Confusion matrix summarizing the cross-validated performance of binary classifiers trained on the outputs of LGN-like model cells to the images in our dataset (compare to Figure 4A). (B) Comparison between the classification accuracy of binary classifiers trained on LGN-like model cells outputs (ordinate) or on spike counts emitted by our 222 SC neurons in the early (40 to 80ms) time window. One data point for each binary classifier (color-coded, see legend). (C) Same as B, but for SC spike counts in the late (90 to 130ms) time window. Error bars denote 95% confidence interval, bootstrapped.

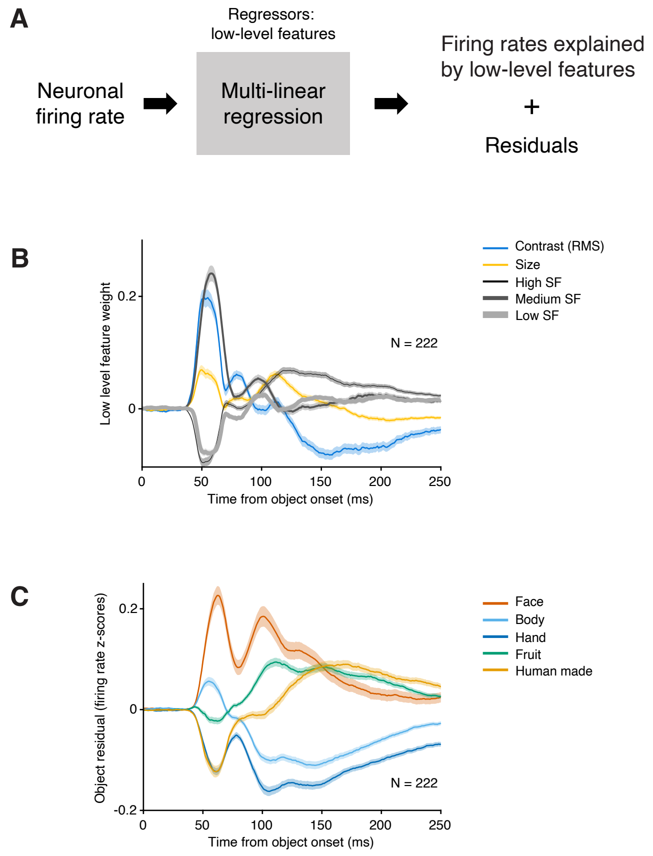


**Fig. S10. Short-latency face preference in visual SC neurons persists even after regressing out the contribution of low-level features.**

(A) Pipeline to evaluate the contribution of five low-level features to the object selectivity we observed in our population of SC neurons. (B) Time-varying weights on each of the five regressors to responses in SC. (C) Residuals split by object category show a strong preference for faces at short latencies, consistent with our other analyses.
